# Predicted pH-dependent stability of SARS-CoV-2 spike protein trimer from interfacial acidic groups

**DOI:** 10.1101/2021.06.06.447235

**Authors:** Vanessa R. Lobo, Jim Warwicker

## Abstract

Transition between receptor binding domain (RBD) up and down forms of the SARS-CoV-2 spike protein trimer is coupled to receptor binding and is one route by which variants can alter viral properties. It is becoming apparent that key roles in the transition are played by pH and a more compact closed form, termed locked. Calculations of pH-dependence are made for a large set of spike trimers, including locked form trimer structures that have recently become available. Several acidic sidechains become sufficiently buried in the locked form to give a predicted pH-dependence in the mild acidic range, with stabilisation of the locked form as pH reduces from 7.5 to 5, consistent with emerging characterisation by cryo-electron microscopy. The calculated pH effects in pre-fusion spike trimers are modulated mainly by aspartic acid residues, rather than the more familiar histidine role at mild acidic pH. These acidic sidechains are generally surface located and weakly interacting when not in a locked conformation. In this model, their replacement (perhaps with asparagine) would remove the pH-dependent destabilisation of locked spike trimer conformations, and increase their recovery at neutral pH. This would provide an alternative or supplement to the insertion of disulphide linkages for stabilising spike protein trimers, with potential relevance for vaccine design.

## 1. Introduction

The spike glycoprotein (S protein) of severe acute respiratory syndrome coronavirus 2 (SARS-CoV-2) has received much attention, as the primary point of attachment between virus and ACE2 receptor on a cell surface [1], and as a target in vaccine design [2]. An original focus on the pre-fusion S protein trimer structure in closed and open states, with receptor binding domain (RBD) down and up, respectively, has expanded to include a further RBD down state, termed locked [3], first observed with in a structure with linoleic acid bound [4]. Here, the terms up and down are used to delineate RBD position, so that the open form maps to RBD up, but both closed and locked forms map to RBD down. This 3 state pre-fusion S trimer picture will be expanded with greater characterisation of post-fusion trimers [5], but the transitions between pre-fusion trimers are of key interest for the early steps in cell entry, and also for the export of fusion competent virus. Further, spike protein trimer expressed at the cell surface is able to mediate fusion between cells [6].

The role of pH has been studied for cell entry and exit of coronaviruses. Both plasma membrane fusion and endosomal mediated fusion pathways can be used [7], with fusion primed by S1/S2 spike protein cleavage during the biosynthetic exit pathway [8]. S1/S2 and S2/S2’ cleavage sites, and the fusion peptide, are critical locations in considering spike trimer conformational transitions [9]. Most cryo–electron microscopy (cryo-EM) structures of spike trimers have been resolved at pH 7.5 – 8. Studies at mild acidic pH have found structural transitions [3, 10], interpreted as relatively favouring closed over locked forms as pH is increased from mild acidic to neutral [3]. Other cryo-EM analysis shows changes in the fraction of RBD up and down conformations between pH values of 6.5 and 8 [11]. Cell biological investigation of the exit pathway for newly synthesised SARs-CoV-2 virus has revealed a novel mechanism where lysosomes are repurposed for viral transit to the cell surface, with a moderate de-acidification of their mean pH from 4.7 to 5.7 [12].

Biophysical and structural analysis of the D614G S protein mutation that emerged in Spring 2020 indicates that it leads to greater availability of the RBD for ACE2 binding, through altered structural transitions [13]. These transitions include packing changes that resemble closed to locked form differences [14]. A model has been proposed in which the locked form is prevalent in the virus export pathway, giving protection of newly synthesised spike proteins at acidic pH, with the D614G mutation coupling to transition between locked and closed conformations [3]. In a complementary model, conformational change in the D614G structure has been proposed to suppress release of S1 from cleaved S1/S2 spike trimers, thus enhancing availability for ACE2 binding [14].

Understanding the factors that stabilise locked, closed, and open S protein trimers is important for vaccine design as well as the pathways of viral infection and biosynthesis. Engineering to stabilise spike trimers [15] is important as knowledge of the most effective conformational targets for vaccine production become apparent [16]. Molecular dynamics simulations are a common tool for structure/function analyses, particular those involving structural transitions. They have been extensively applied to SARS-CoV-2 S protein [17], including studies of receptor binding [18], drug binding to the linoleic acid binding pocket [19], the opening of cryptic pockets as potential drug targets [20], temperature effects [21], and the effect of glycosylation [22].

Specifically for pH-dependent effects, several computational methods are available. Constant pH molecular dynamics has been applied to coronavirus protease active sites, reporting on the protonation states that may be best suited to inhibitor design [23]. Atomistic simulations in general are mostly low throughput in terms of the starting coordinate sets that are studied. At the commencement of the current study, about 130 structures were available for SARS-CoV-2 S protein trimers. Since these represent substantial variability, for example locked, closed or open forms, D614 or G614, neutral pH or acidic pH, high throughput computational methods have a role to play, in this case to increase understanding of pH-dependent properties. Based on earlier work [24], a web server for p*K*_a_ calculations has been benchmarked against known key sites for the acidic pH-dependence of influenza virus hemagglutinin [25]. This work was extended with a dataset of 24 spike trimers, collated in late Summer 2020 and mostly lacking what is now recognised as representatives of the locked form. A sparse set of groups predicted to couple with pH-dependence of closed to open form transition was found, focusing on the environment around a single salt-bridge [20].

The locked form is now more extensively characterised, with about 10 S protein trimers at the end of January 2021, as well as structures at mild acidic pH, and with the D614G mutation. Accordingly, the focus shifts to the transition between closed and locked forms in the current report, revealing a wider set of acidic groups (including D614) that are predicted to modulate stability at mild acidic pH. These groups are at interfaces that tighten in the locked form, relative to the closed form. Indeed, the change in burial of monomers within the S protein trimer is greater for the closed to locked transition, than for the closed to open transition. Predicted destabilisation in the locked form at neutral pH can be offset by either transition to the closed form or reduction to acidic pH. Alternatively, mutation of these acidic groups is predicted to stabilise locked form trimers at neutral pH.

## 2. Methods

### 2.1 Structural data

A set of 129 SARS-CoV-2 spike trimer structures was retrieved from the RCSB/PDB [26] in January 2021, from a list of list generated by probing the SARS-CoV-2 spike sequence, UniProt [27] id P0DTC2, with the Basic Local Alignment Search Tool (BLAST) [28] into the RCSB/PDB at the National Center for Biotechnology Information, and subsequent checking for SARS-CoV-2 amongst the coronavirus spike proteins returned. Non-spike binding partners were removed, leaving the 129 trimers and 387 monomers (spike387). The monomers of spike387 are listed in Supplementary Table 1, along with assignment of RBD up or down according to PDB entry, and confirmed by visual inspection. Subsequent to analysis of spike387, sub-groups of locked, closed, and open S protein trimers were made, each trimer being uniformly RBD down (locked or closed), or RBD up (open). To maintain balance, 8 trimers were used for each sub-group, again with binding partners (ACE2, antibody fragments) removed. Further sub-groups were formed, consisting of: a single trimer with an engineered disulphide link (6zoz [29], termed diSlocked), 3 trimers at acidic pH [10] (pHlocked), and 8 trimers carrying the D614G mutation (D614Gset). Constituent monomers within trimers of the 6 sub-groups, along with RBD up or down designation, are given in Supplementary Table 2.

### 2.2 Sequence data

Sequences of spike proteins from the seven coronaviruses currently known to infect humans were aligned with the Clustal Omega [30] implementation at the European Bioinformatics Institute [31], and amino acid conservation relative to SARS-CoV-2 spike protein noted at particular sites. The seven S protein sequences included (with UniProt identifiers) were SARS-CoV-2 (P0DTC2), SARS (P59594), MERS (K9N5Q8), HCoV-HKU1 (Q5MQD0), HCoV-OC43 (P36334), HCoV-NL63 (Q6Q1S2), and HCoV-229E (P15423).

### 2.3 Structure-based calculations

Solvent accessible surface area (abbreviated to ASA) was calculated for S protein monomer burial within a trimer. For each monomer within a trimer (after non-spike components have been removed), ASA was calculated for the monomer alone and for the monomer within the trimer, the difference (d-ASA) giving the extent of burial. The contribution of polar and non-polar atoms to this burial was recorded, following previous methodology [32]. To facilitate comparison between sub-groups, d-ASA values were averaged over monomers within a sub-group. A key feature is that all calculations were referenced to an alignment of amino acid sequences for all monomers (spike387 and additional monomers added in subsequent formation of sub-groups). This ensures that all comparisons and heat maps are correctly aligned, allowing for engineered monomers and residues present in a sequence but disordered in a structure. Amino acid numbering throughout is given as the equivalent location in UniProt entry P0DTC2. Missing coordinates are not modelled, structure-based calculations use only the ordered residues reported in each structure. From vectors of 1 (ordered) and 0 (disordered) running along the sequence of each monomer, similarity to chain A of the locked conformation 6zb4 was calculated by vector dot product. Monomers were then listed in heat maps (of order/disorder or d-ASA) according to similarity with 6zb4A.

Electrostatics calculations used p*K*_a_ predictions following the Finite Difference Poisson Boltzmann (FDPB) / Debye-Hückel (DH) hybrid method, termed FD/DH [24]. Ionisable groups that are not buried can sample both FDPB and DH schemes, and these calculations are combined with Boltzmann weighting. Since the latter is a simple water-dominated and highly damped estimate of electrostatic interactions, non-buried groups will not have high p*K*_a_ deviations (Δp*K*_a_s) from normal values, unless those are stabilising. On the other hand, buried groups, shielded from water-dominated interactions, can have Δp*K*_a_ values that relate to substantial destabilisation as well as stabilisation. The method has been implemented in a web tool, following testing with sites known to determine the pH-dependence of influenza virus hemagglutinin stability [25]. Here, the focus is on predicted stability change between pH 7.5 (extracellular, cytoplasmic pH) and the lower pH experienced in import and export pathways (pH 5 was used as the lower value). This pH-dependent energy can be derived using the relationship [33] *ΔG_pH-dep_ = ∫(2.303RTΔQ)dpH*, where ΔG_pH-dep_ is the pH-dependent contribution to conformational stability (over the pH range of the integration, 7.5 to 5), differenced between two states. The states here are ionisable groups interacting in the FD/DH scheme, and the same set without interactions (null state with normal model compound p*K*_a_s). ΔQ is the charge difference between those two states, R is the Universal Gas constant, and T is set at 298.15 K. Since the null state is uniform between structures, results for each structure can be compared to assess differential responses to pH change in the mild acidic range. Groups responsible for the predicted effects can be identified since ΔQ separates into component ionisations.

Molecular viewers used to analyse spike proteins were NGL [34], Swiss PDB Viewer [35], and PyMOL. For representative trimer structures in the locked (6zp2), closed (6zp1) and open (7a98) sub-groups, d-ASA values were transferred to the B factor column (capped at 99 Å2), facilitating visualisation of monomer burial within a trimer.

## 3. Results and Discussion

### 3.1 Differences in monomer burial between S protein trimer forms are more extensive than are order-disorder transitions

Monomers within the spike387 set were classed as RBD up or down, noting that down does not distinguish between closed and locked trimers. Clustering of monomers was undertaken according to similarity of vectors representing order/disorder, and referenced to the A chain monomer within the 6zb4 locked trimer [4]. The resulting heat map of order/disorder is shown alongside a record of up or down RBD, and S protein domain structure (Fig. 1). Most clear-cut is ordering around position 840 in a set of structures with down RBDs and known to be in the locked form, at the top of the heat map. Other clusters of monomers, in terms of order/disorder, do not uniformly map to up or down RBD.

**Fig. 1.**
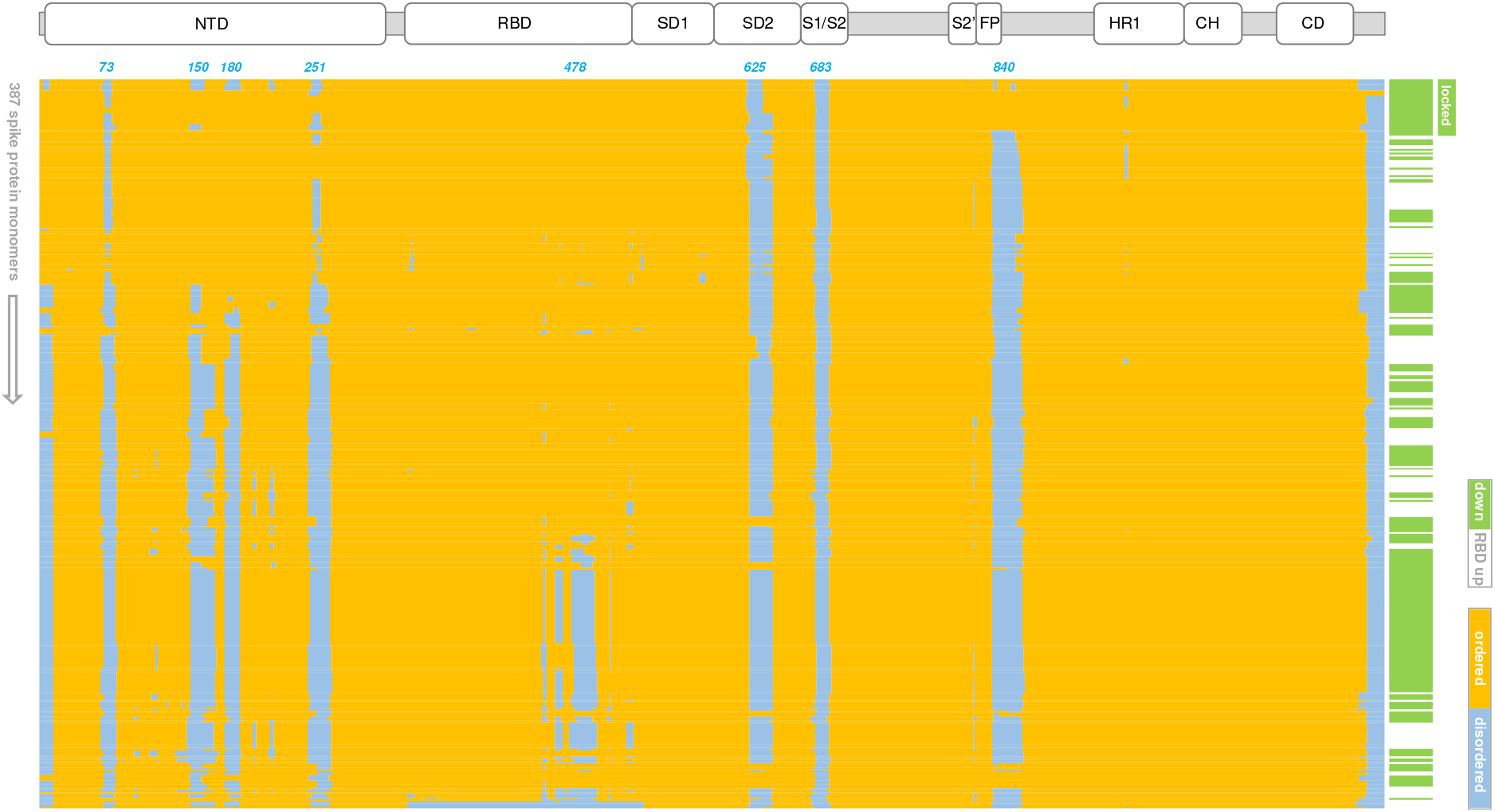
Spike protein monomers clustered according to similarity of ordered/disordered regions. The spike387 set is listed vertically according to similarity with chain A of 6zb4 (top). Order (yellow) and disorder (blue) are plotted. A smaller vertical heat map indicates monomers with down (green) or up (white) RBD positions. A group of locked conformations that cluster with 6zb4 monomers is indicated. Amino acid sequence (horizontal) covers the extent of ordered regions in spike ectodomain structures, with specific regions of disorder noted along the top of the heat map. Domain structure of the spike protein is also shown, in register with the heat map.

A similar procedure was used to map monomer burial within trimers. For each monomer, ASA loss upon integration of that monomer within the trimer was calculated (d-ASA). The order of monomers in the d-ASA heat map of Supplementary Fig. S1 is the same as that in Fig. 1. Summed d-ASA for each monomer is also shown. Known locked conformations mostly exhibit higher d-ASA than other conformations, including closed. The region around 840 in the d-ASA heat map differentiates the locked form again, but there is also a greater richness in differences across the sequence, particular for the RBD, reflecting a tightening of interfaces in the locked form relative to closed form. This analysis gives a view of detailed changes that complement molecular graphics comparison of locked and closed RBD down structures.

### 3.2 Relatively polar interfacial interactions differentiate locked from closed trimers

In order to more easily compare different conformational forms of spike trimer, sub-groups from spike387 were made with either all RBD down or all RBD up monomers. Three sub-groups (locked, closed, open) were matched numerically at 8 trimers (24 monomers) each. Other sub-groups were added: a single locked conformation trimer with engineered cysteines creating a disulphide link [29] (diSlocked), 8 structures with the D614G mutation (D614Gset), and 3 structures at acidic pH values (pHlocked). For D614Gset, trimers have various combinations of up and down RBD containing monomers. Summed d-ASA values show clear distinction of locked and diSlocked forms from closed, and open forms (Fig. 2a). The change of monomer burial within a trimer from locked to closed is greater than that between closed and open, emphasising the interfacial tightening of the locked form. These changes are widespread and not solely located in regions that undergo disorder to order transitions (Fig. 1). Monomers of the pHlocked set have d-ASA intermediate between locked and closed trimers, whilst D614Gset monomer d-ASA values span a much greater range than other sub-groups.

**Fig. 2.**
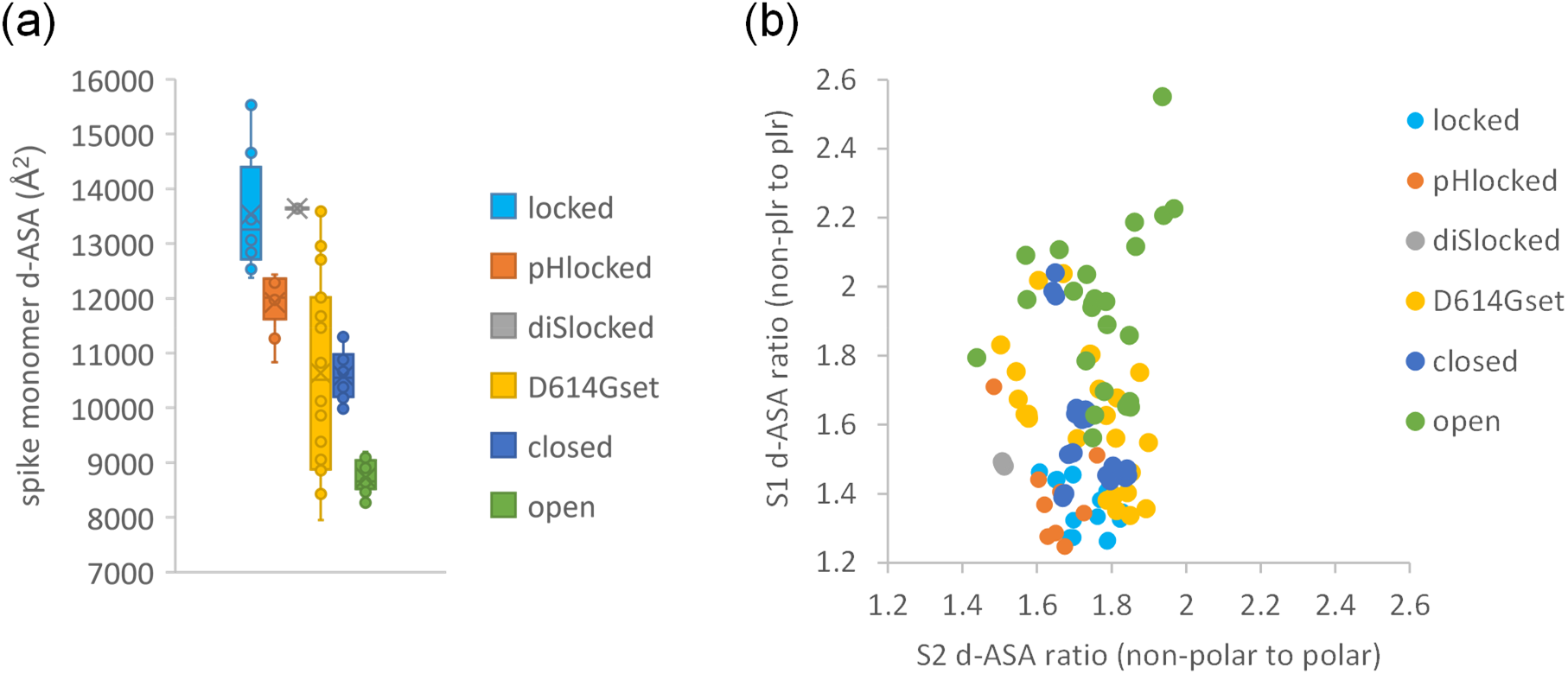
Buried ASA of monomers within spike trimer, compared between sub-groups. (a) A scatter plot showing the ratio of non-polar to polar buried d-ASA for the S1 and S2 subunits, upon monomer burial in the trimer. Variation of this ratio is much greater for S1 than for S2 (seen with equal scaling of the axes), with higher values for the open form indicating that residual buried surface with RBD up is relatively non-polar. (b) Box and whisker plot for d-ASA distributions within each sub-group.

The ratio of non-polar to polar contributions to d-ASA was calculated for the S1 and S2 subunits of each monomer (Fig. 2b). For external protein surfaces not involved in protein-protein interactions the most common value of this ratio, in a distribution over patches, is about 1.2, increasing for more hydrophobic, interacting surfaces [32]. For the open sub-group, with least monomer burial, non-polarity increases for both S1 and S2 contributions. Conversely, monomer interface within a trimer becomes successively more polar for sub-groups that have higher d-ASA, with polarity for S1 d-ASA in the locked form close to that seen for non-interfacial surface (Fig. 2b). It can be concluded that transition from RBD up to RBD down, and further transition to the locked form, involves burial of surface that is not especially hydrophobic.

### 3.3 Acidic sidechains at tightened interfaces are predicted to have elevated pK_a_s

Comparison of Fig. 3a and Fig. 3b reinforces the observation that d-ASA differences between S trimer sub-groups are more extensively spread over spike monomers than are order/disorder differences. The extent of d-ASA change between locked and closed forms, in comparison with that between closed and open forms, is also emphasised. In light of reported pH-dependent transition between S trimer conformations [3, 10], p*K*_a_ predictions were made for ionisable residues in trimer sub-groups. To directly address changes in the mild acidic pH range, the predicted contribution to change in conformational stability between pH 7.5 (cytoplasm, extracellular) and pH 5 (endosome) was calculated (summed and for each ionisable group). The results are added to Fig. 3 as a heat map of calculated per-residue pH-dependence, averaged over monomers within each sub-group (Fig. 3c). Towards the C-terminal part of S2 are several histidines [25, 36], for which burial and predicted resistance to protonation from pH 7.5 to 5 leads to destabilisation in all sub-groups. Of more interest for differences between sub-groups are amino acids with acidic sidechains that have elevated p*K*_a_s and are predicted to be stabilising from pH 7.5 to 5, since destabilisation is reduced. Destabilisation derives from a local environment that suppresses ionisation at a pH above the normal p*K*_a_. Several Asp and Glu sidechains are of interest, particularly those for which the effect is exhibited in locked forms but not others, as a consequence of the increased burial.

**Fig. 3.**
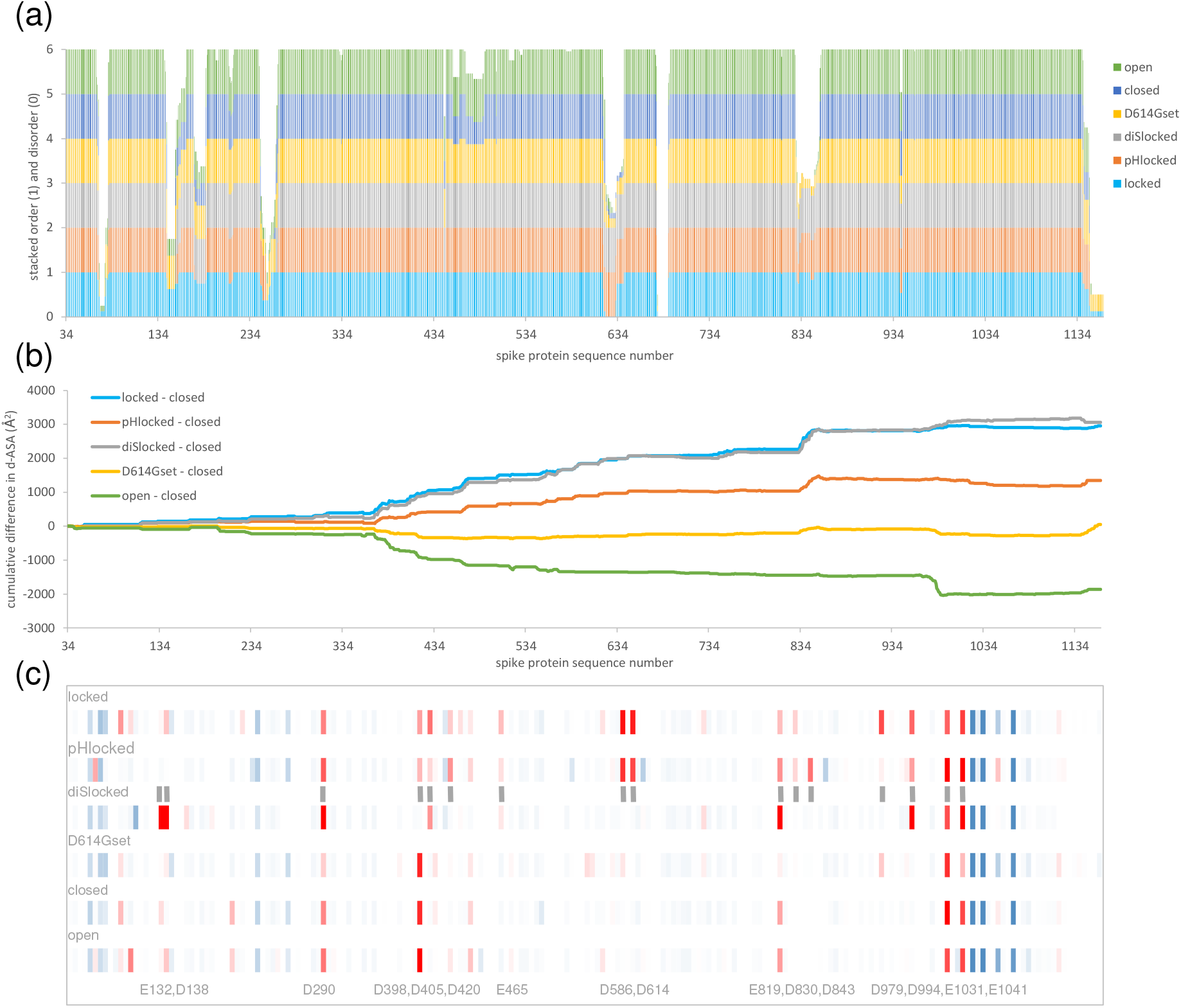
Acidic groups with increased burial in the locked form are predicted to give a mild acidic pH-dependence. (a) Order/disorder is illustrated with stacked histograms (1/0) for the 6 sub-groups of spike trimers. Differences of d-ASA between sub-groups are more extensive than are those of order/disorder, illustrated with cumulative plots of the closed form subtracted from the 5 other sub-groups (b). The magnitude of locked to closed difference is larger than that of open to closed. (c) A heat map of predicted conformational stability change from pH 7.5 to 5, for each sub-group (labelled), with stabilising (red) or destabilising (blue) indicated. Amino acids that differ the most between sub-groups are indicated at the bottom of the plot, together with additional grey vertical bars aligned with the heat map. Panels (a) and (b) are in register. Panel (c), with only ionisable groups, is approximately in register with the full sequence plots.

### 3.4 Predicted stabilisation of locked forms, and destabilisation of closed and open forms, from pH 7.5 to pH 5

Summed values of predicted pH-dependence give a stabilisation of locked form sub-groups as pH falls from 7.5 to 5, but a destabilisation of other sub-groups over the same range (Fig. 4). The consequence of these predictions is a model in which the locked form is preferentially stabilised in low pH secretory pathway vesicles, whereas closed (unlocked) and open forms are stabilised at extracellular, neutral pH, as suggested from structural studies [3].

**Fig. 4.**
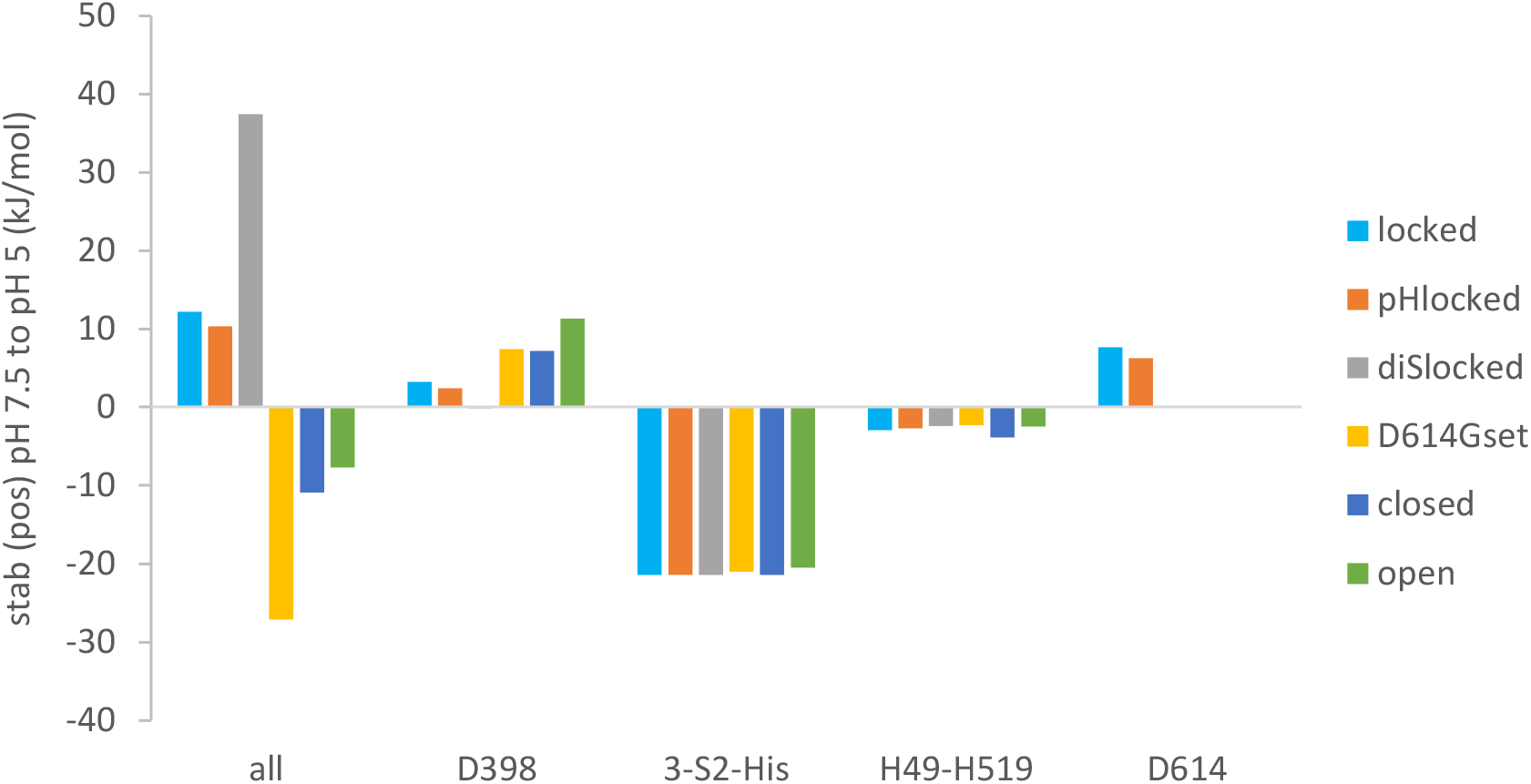
Predicted pH-dependent stabilisation for different conformational forms. Calculated stabilisation or destabilisation (pH 7.5 to 5) is shown for the average over each sub-group of: all ionisable groups, D398, H1048 + H1064 + H1088 of S2, H49 + H519, D614.

Looking at individual group contributions, 3 His in S2 (H1048, H1064, H1088) are buried and largely inaccessible to solvent throughout the sub-groups, so that they give a uniform destabilisation over the pH 7.5 to 5 range, as each His transits through the p*K*_a_ (6.3) that would normally mark ionisation. These groups are not predicted to contribute differentially to pH-dependence of locked, closed, and open forms. Next, D398 was proposed to couple to pH-dependence of closed (mostly unlocked) to open forms [36]. This result is replicated in the current larger study (Fig. 4), with a predicted D398 contribution to variation of pH-dependence between locked, closed, and open forms, but in the opposite sense to the overall effect. Two histidines have been suggested as potential mediators of locked trimer stabilisation at acidic pH [3], but no clear differential contributions of H49 and H519 are seen between sub-groups of structures in the current study (Fig. 4).

### 3.5 D614G mutation and structural preferences

A feature of predicted pH-dependence for which the picture changes when locked (RBD down) structures are separated from closed (RBD down), is the influence of D614. Contributions of D614 to the calculated pH-dependence are seen for locked and pHlocked sub-groups, in a sense to stabilise those forms at pH 5 relative to pH 7.5 (Fig. 4). It has been reported that the S protein D614G mutation increases virus infectivity [37], and suggested that the mutation leads to a greater population of up relative to down RBDs due to loss of an inter-monomer latch formed by the sidechain of D614 [13]. Recent structural studies with an S protein ectodomain engineered with D614G, report that the increase in infectivity results from stabilisation of the S trimer against dissociation [5]. One of the structures (7krq) from that D614G study has all RBDs down, and d-ASA for monomers in this structure (Supplementary Fig. S2) are similar to those seen for locked forms (Fig. 2a), consistent with trimer stabilisation. Generally, the D614Gset sub-group exhibits a wide distribution of monomer d-ASA values, but these structures mostly have mixed up and down RBD within each trimer (Supplementary Fig. S2).

D614 is predicted to be relatively destabilising at neutral pH in locked and pHlocked forms (Fig. 3c, Fig. 4). Even after incorporating stabilising D614 sidechain interaction with K854, the reduction in solvent exposure due to ordering of the loop around residue 840 leads to a predicted average destabilisation, at pH 7.5 compared to pH 5 and averaged over the locked sub-group, of 7.7 kJ/mol. This implies a mean D614 p*K*_a_ shift to about 6.0, from the intrinsic value of 4.0, consistent with wide-spread D614 - K854 salt-bridge formation in locked structures at pH 7.5 or 8. There are examples of this salt-bridge in all sub-groups (other than D614Gset), although at lower incidence outside of the locked sub-group. Our interpretation is that D614 - K854 interaction is solvent exposed and relatively weak outside of locked foms. Then, ordering of the loop around residue 840 in the locked form decreases solvent exposure, putting a greater onus on salt-bridge formation to reduce destabilisation of D614. Summed over interactions though, destabilisation of D614 is predicted to result for the locked form, and will be greater at pH 7.5 than at pH 5. This model is consistent with stabilisation of the D614G S trimer against dissociation at neutral pH [38]. Referring to the 3 structures in the pHlocked sub-group, for two (6xlu, pH 4.0 and 7jwy, pH 4.5) the 840 loop is ordered. In both of these, K854 is outside of salt-bridge range, consistent with elevated p*K*_a_ and protonation of D614 at acidic pH.

### 3.6 Predicted locations of pH-dependence flank regions of known structural sensitivity

In common with D614, other predicted sites of pH-dependence are coupled to a tightening of monomer-monomer interfaces in the locked form S trimer (Fig. 5a), so that solvent exposure is reduced and Asp/Glu sidechain p*K*_a_s are elevated, with predicted destabilisation of the locked form as pH increases through the mild acidic pH range. Presumably, pH-independent features, such as linoleic acid binding [4], also couple to the energetic balance between locked and closed forms. For the pH-dependent component, it is notable that a tendency towards a locked form at acidic pH is still evident for the D614G S protein trimer [3], consistent with the existence of other pH-dependent sites (Fig. 3c, Fig. 5a). Whereas the D614 - K854 interaction is between monomers, neighbouring basic residues for other acidic sites of predicted pH-dependence, where they exist, tend to be from the same monomer. Nevertheless, the predicted effect is the same, it is the solvent exposure of the acidic group rather than the details of salt-bridging that is the key determinant of predicted pH-dependence. Several of the implicated acidic residues are present in only SARS-CoV-2 and SARS, of coronaviruses known to infect humans (Fig. 5b).

**Fig. 5.**
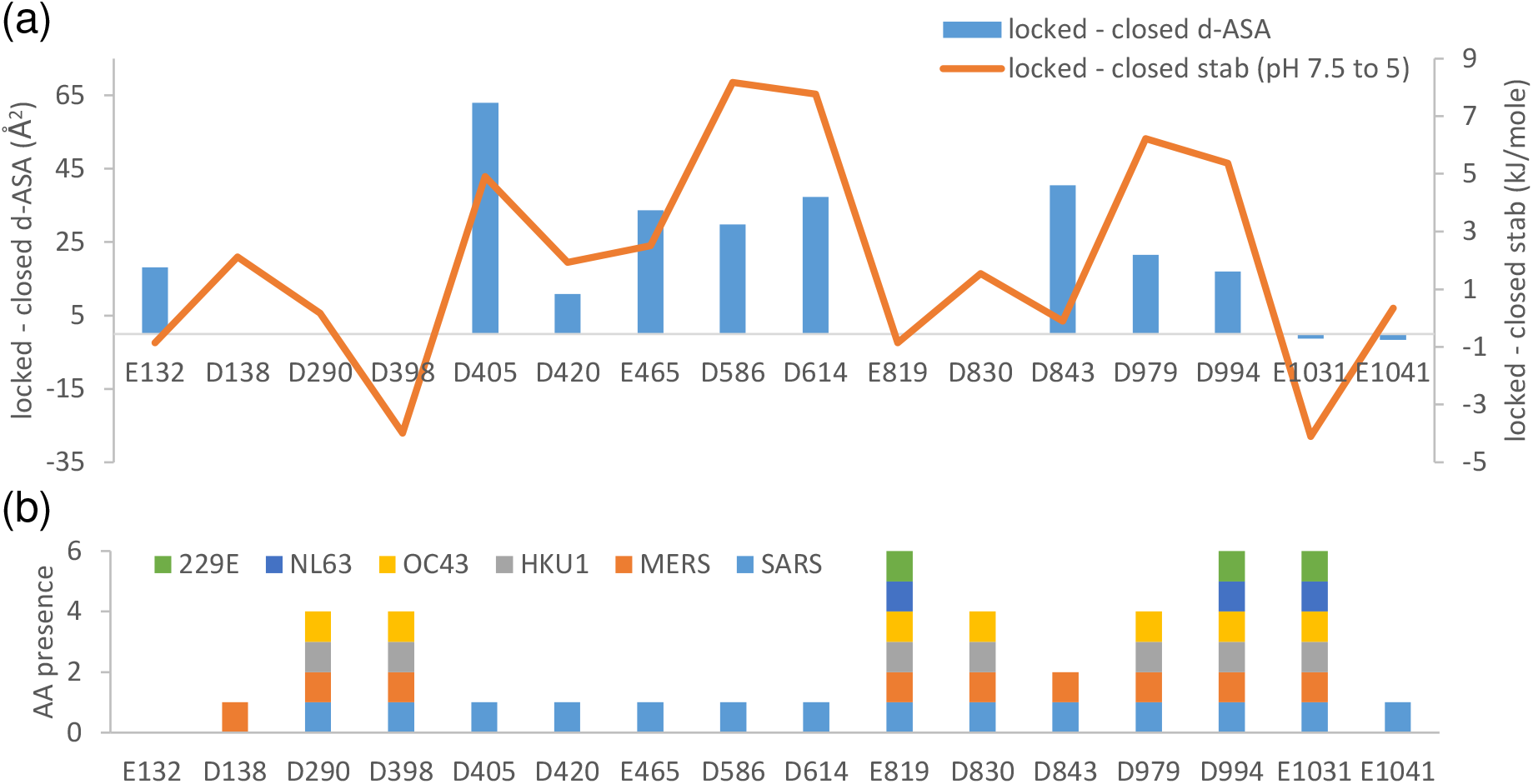
Individual residue contributions to locked / closed form differences. (a) For amino acids highlighted in Fig. 3c, both d-ASA and predicted pH 7.5 to 5 stability change are plotted as differences (locked – closed). (b) The presence of these residues in a set of coronaviruses that infect humans (other than SARS-CoV-2) is shown with a stacked histogram. For example, E132 is present only SARS-CoV-2, whilst D994 is also present in the other 6 viruses.

Display of d-ASA on S protein structure shows the expected difference between open (Fig. 6a) and closed (Fig. 6b) forms, and emphasises the difference (that is not evident by overall RBD location) between closed (Fig. 6b) and locked (Fig. 6c) forms. General location (Fig. 6d) and detail (Fig. 6e) are shown for residues that are present in the more compact interfaces of the locked form trimer, and which are predicted to contribute to pH-dependence over the mild acidic range. Some residues (D586, D614, D830, D843) flank regions of known importance for structural transitions (S1/S2 and S2/S2’ cleavage sites, fusion peptide), and other residues (D405, D420, E465, D979, D994) are adjacent to a site commonly engineered to generate more stable trimers [3]. Additionally, sites of predicted pH-dependence in the RBD are proximal to the linoleic acid binding site [4]. It is therefore suggested that members of this set of acidic residues (Fig. 5b, Fig. 6e), beyond D614, couple pH-dependence to conformational biases of functional relevance.

**Fig. 6.**
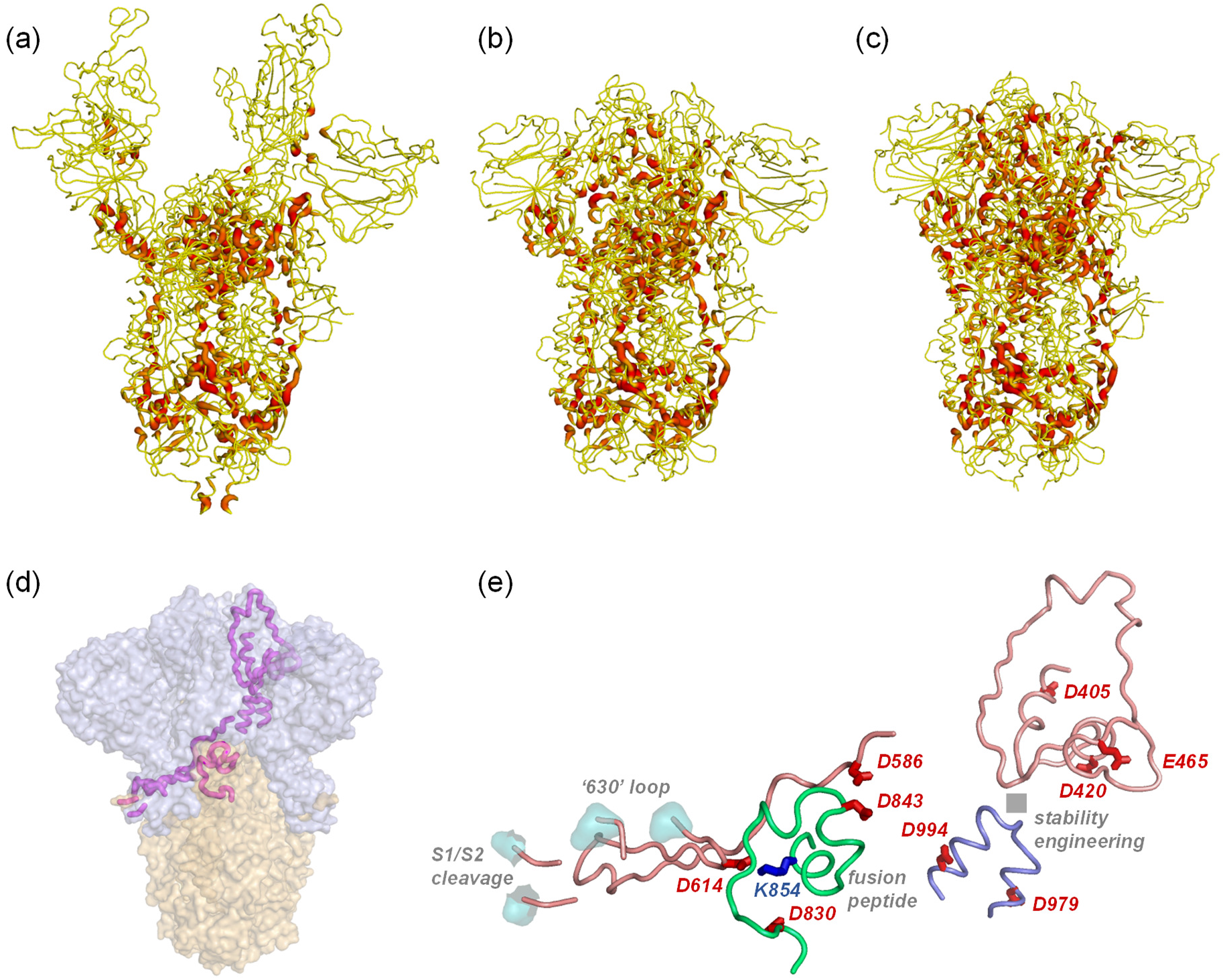
Location of sites predicted to differentiate the mild acidic pH-dependence of locked and closed forms. Tube plots of spike protein backbone are shown with colour-coding from d-ASA of 0 Å^2^ (yellow) to 80 Å^2^ (red) for open/7a98 [9] (a), closed/6zp1 [29] (b), and locked/6zp2 [29] (c) forms. Within the regions of increased d-ASA in locked relative to closed forms are amino acids with acidic sidechains, shown in a coarse view with magenta tube plot (one of 3 symmetry-related representatives) on a background of spike trimer S1 (grey) and S2 (orange) subunits (d). This region is drawn in greater detail (e) with backbone tube and amino acid stick plots, noting that this one of the 3 representatives consists of regions from all 3 monomers (colour-coded tube). Sites of interest with regard to conformational change and stability within the trimer are also indicated.

### 3.7 pH-dependence generated through acidic groups at interfaces

In this model for pH-dependence of locked to closed form transition, a surface acidic group, whether or not paired with a basic group, loses substantial ASA on formation or tightening of an interface. The acidic p*K*_a_ increases in the range pH 4 to 7. A bias against interface formation is introduced, which is relieved as pH reduced. Depending on the conformational details and interactions with neighbouring groups, a buried acidic group could be stabilising, but that is not predicted here.

Although functional ionisation of carboxylate sidechain groups at mild acidic pH is most well-known for acid-base catalytic systems, such as the elevated p*K*_a_ of E35 in hen egg white lysozyme [39], many non-catalytic occurrences have been discussed. In viruses, coiled-coil formation in a variant influenza hemagglutinin is pH-dependent between pH 4.5 and 7.1, possibly mediated by E69 and E74 [40]. A model for pH-dependent association of some antigenic peptides and class II proteins of the major histocompatibility complex suggests that buried interfacial Asp or Glu residues mediate this process, with p*K*_a_s higher than 7 [41]. Interfacial residues D612 and E613 of PsaB in photosystem I are proposed to be responsible for mild acidic pH-dependence of electron transfer in complexes with plastocyanin or cytochrome *c*_6_ [42]. The periplasmic chaperone HdeA of *Escherichia coli* acts on a shift from neutral to lower pH *via* dimer dissociation and partial unfolding, these changes mediated by two aspartic acid residues [43]. In a further example, aspartic acid residues have been proposed as determining neutral to low pH conformational variation in an antigen-binding region of a therapeutic monoclonal antibody [44].

Given the background of acidic sidechains mediating mild acidic pH-dependent effects, it is reasonable to investigate similar interactions when considering the pH-dependence of spike protein conformation, supplementing the assessment of histidine protonation in regard to pH-induced changes in viruses [45].

## 4. Conclusions

Analysis of spike trimer structures shows the expected large ASA change, for monomer burial, between closed and open forms, and an even larger change and interface tightening between closed and locked forms. A subset of acidic groups was identified in the locked form monomer-monomer interfaces that are predicted to mediate pH-dependent stability at mild acidic pH. These are located in regions that flank sites of known importance for stability engineering and/or function, such as cleavage sites and fusion peptide. D614 is amongst the groups identified and, in common with other acidic groups in the subset, is predicted to destabilise the locked form at neutral pH. This result is consistent with a report that the G614 mutation can exist in a form that resembles locked and stabilises the trimer at neutral pH, leading to suppression of S1 subunit loss and increased RBD – ACE2 binding [38].

The current work is also consistent with a model for pH-dependent conformational masking of RBDs within the S protein trimer, derived from the 3 structures of the pHlocked set [10]. That study highlighted D614 and D586 (in common with our calculations), and also D574. If the prediction of a wider set of acidic groups relevant for pH-dependence is correct, then it might be asked why, from this set, only mutation at D614 has been seen widely seen in SARS-CoV-2 genomes. Of note though, a study of recurring spike protein mutations highlights several regions that overlap with, or are adjacent to, segments identified in Fig. 6e [46]. These mutations could influence folding and packing around the sites predicted to mediate pH-dependence. Since pH modulation of the D614G spike trimer conformation has been reported [3], there must be groups beyond D614 responsible for some degree of pH-coupling.

The recently discovered lysosomal exit pathway for newly synthesised SARs-CoV-2 virus provides a mild acidic environment, even including the observed moderate de-acidification [12]. Reduced acidification is thought to reduce activity of lysosomal proteases, thereby protecting exposed viral proteins from proteolysis, beyond the S1/S2 priming cleavage of S protein that confers an advantage for SARS-CoV-2 infection [47]. Stabilising the locked form at acidic pH is proposed to contribute to protection of the spike protein along the secretory pathway [3]. The current results are consistent with this model, extending the set of groups that may be responsible for pH-dependent stabilisation (beyond D614). Importantly, the balance between priming cleavage and stabilisation along the biosynthetic pathway is the focus for ongoing viral adaptation, including the observation that P681R mutation in the B.1.617 lineage optimises the furin cleavage site and could enhance transmissibility and pathogenicity [48].

An implication of the current work is that it should be possible to engineer more stable locked forms at neutral pH through mutation of the acidic groups, (e.g. Asp to Asn), at sites indicated in Fig. 6e. In line with the observation of close packing around G614, Asp to Gly changes could be tested, although the D614N mutation also leads to a more compact spike trimer [49]. Such analysis would probe the balance of stabilising and destabilising contributions to spike trimer structure during trafficking, and it could provide further insight into production of specific spike protein conformational forms for vaccines.

## Acknowledgements

The authors thank Lorena Zuzic for discussion, and the computational shared facility at the University of Manchester for support and resources.

---

**Supplementary Table 1.**
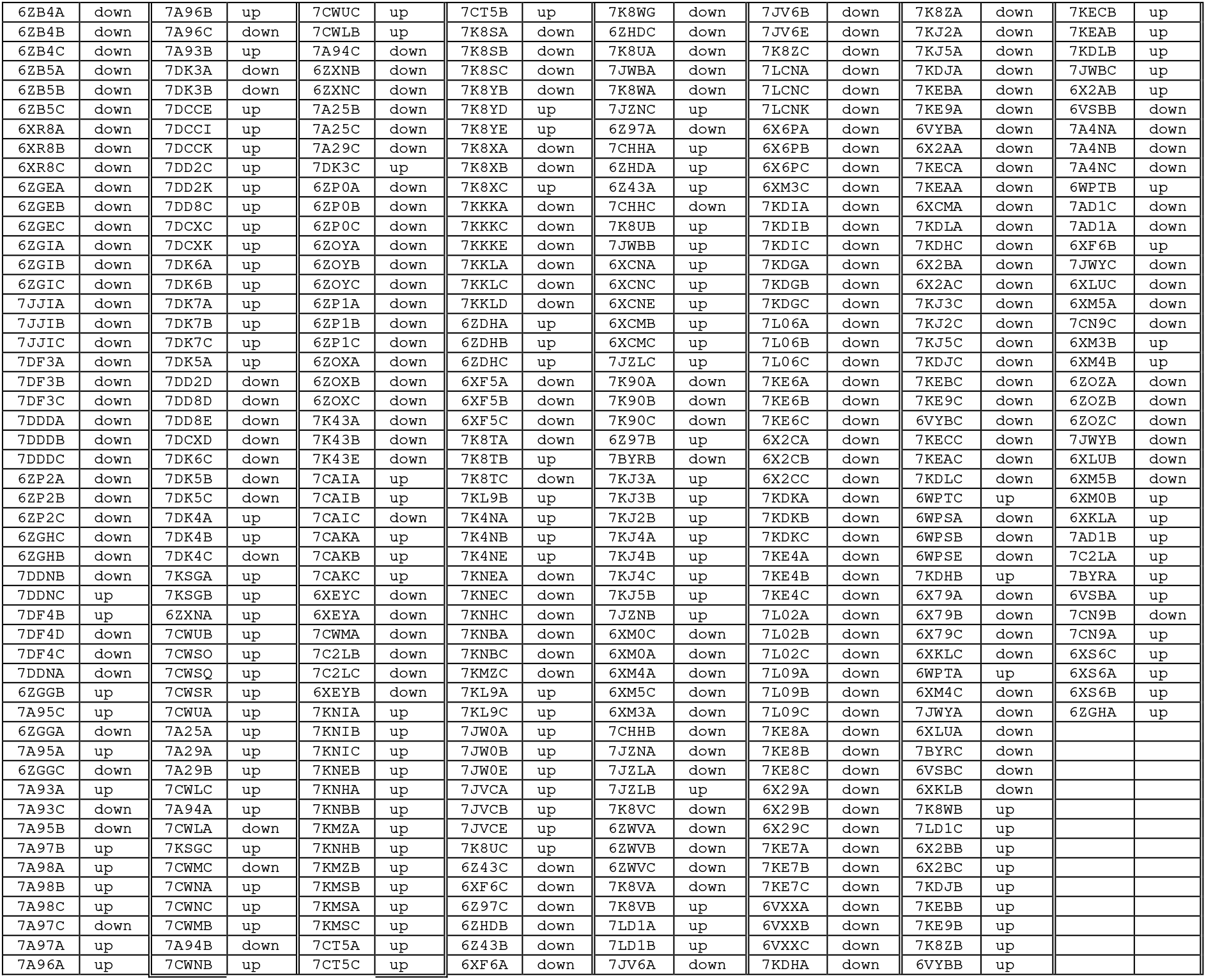
Monomer PDB/chain identifiers are given in pairs with down or up states for the RBD. These 387 monomers from 129 spike protein trimers form the spike387 set. The order of monomers (left to right, top to bottom) is that given in the clustered heat maps of Fig. 1 and Supplementary Fig. 1.

**Supplementary Table 2.**
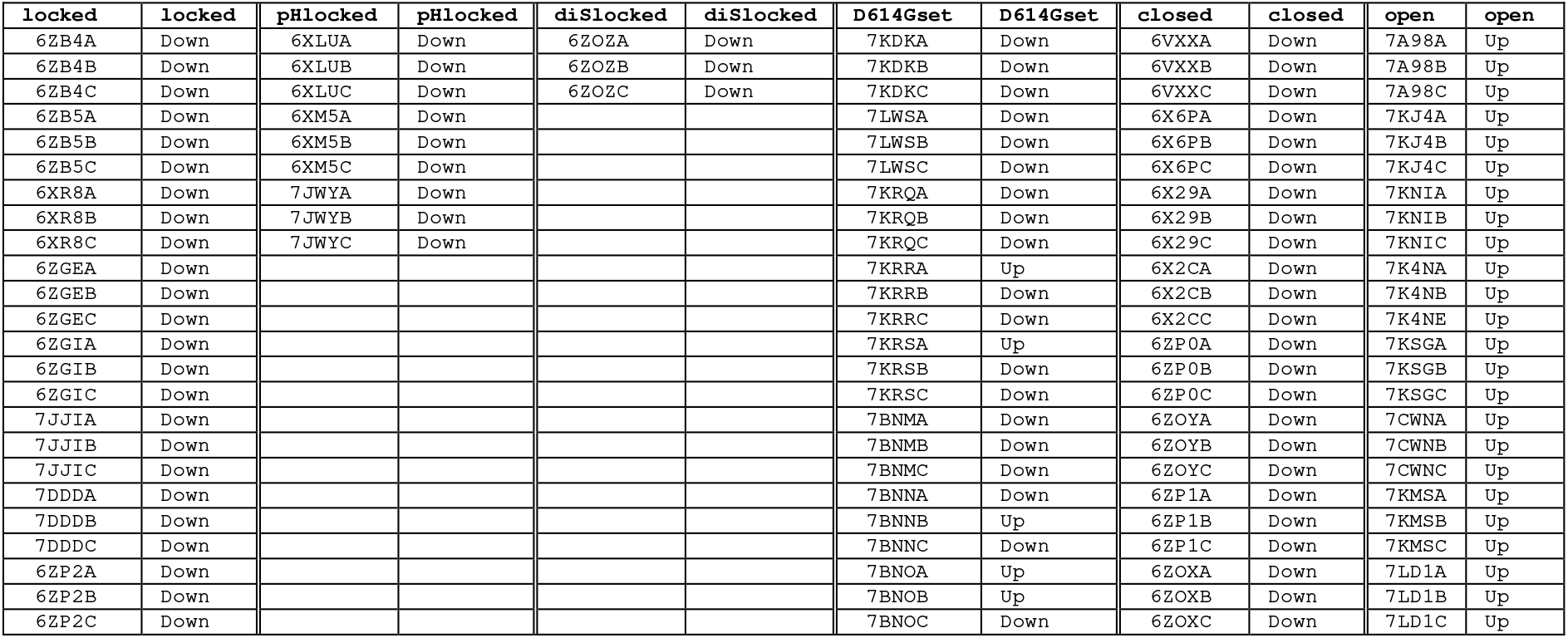
Monomer PDB/chain identifiers are given in pairs with down or up states for the RBD, S protein trimers in each of 6 sub-groups (labelled by the column headers).

**Supp. Fig. 1.**
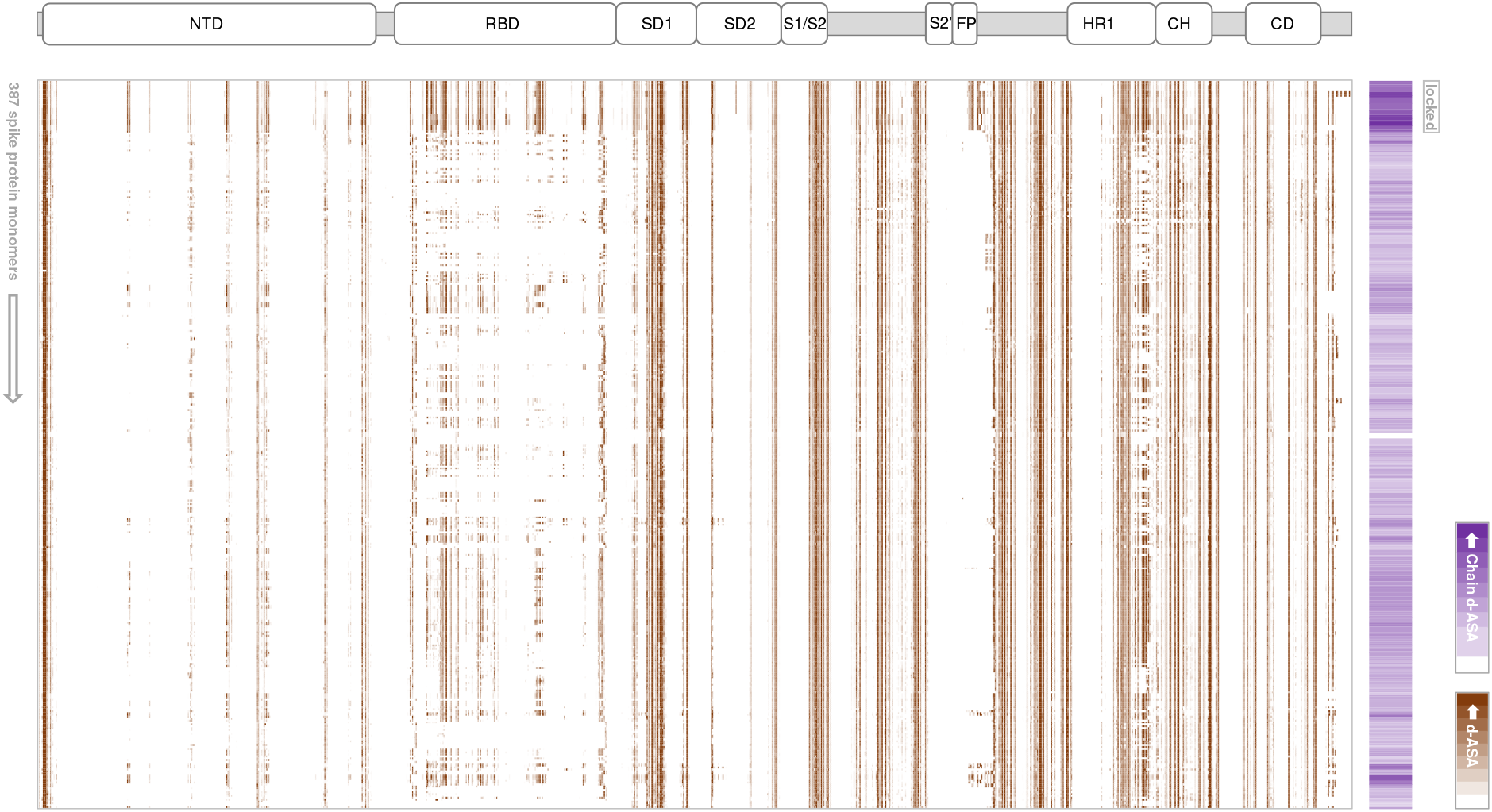
Spike protein monomer burial within a trimer. Monomers are listed vertically according to the clustering derived for Fig. 1 (6zb4A and a group of locked forms at the top), and with sequence variation horizontally, referenced to the domain structure. A heat map of burial ASA (d-ASA) for monomer within trimer is shown, from zero (white) with increasing (brown) colour, capped at d-ASA = 80 Å^2^. A smaller vertical heat map gives d-ASA per monomer (white, smaller to purple, greater).

**Supp. Fig. 2.**
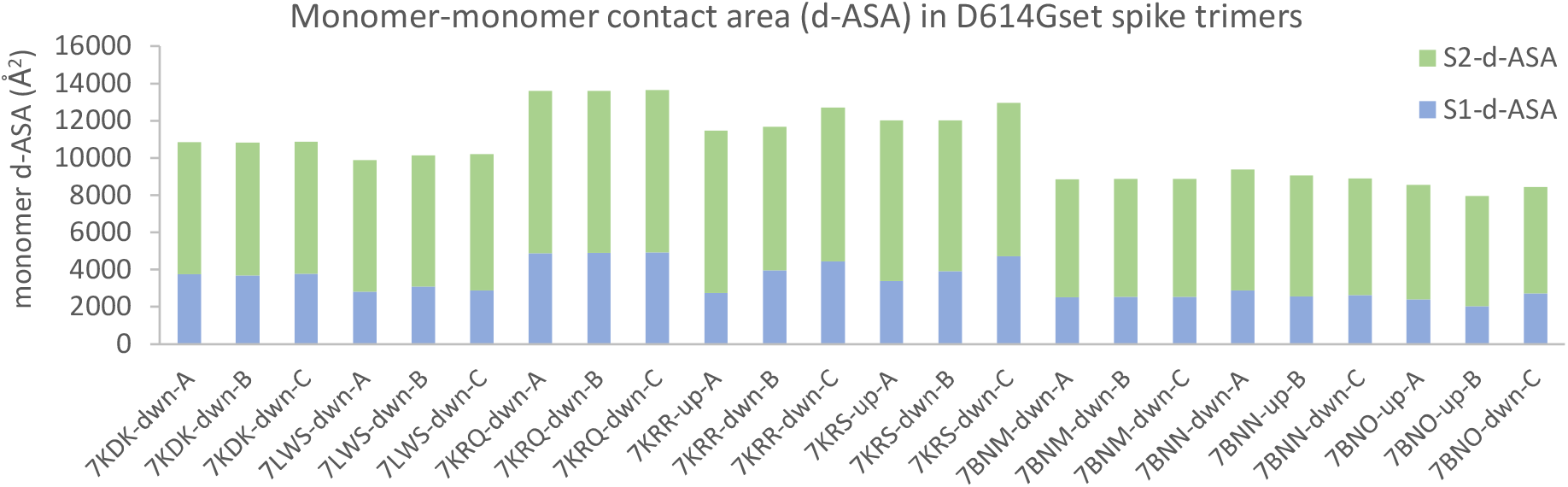
Monomer burial within trimer for members of D614Gset. Variation in both S1 and S2 subunit contributions to d-ASA contribute to the large overall range of d-ASA values exhibited within D614Gset. RBD up or down monomer conformations are indicated.

